# MetaTracer: A nucleotide alignment-based framework for high-resolution taxonomic and transcript assignment in metatranscriptomic data

**DOI:** 10.64898/2026.02.20.707109

**Authors:** Tara N. Furstenau, Isaac Shaffer, Kuei-Ling C. Hsu, Talima Pearson, Robert K. Ernst, Viacheslav Y. Fofanov

## Abstract

**Summary:** MetaTracer is a nucleotide alignment-based tool for metatranscriptomic analysis of complex bacterial communities that assigns sequence reads to both taxonomic groups and expressed genes in a single pass. Full nucleotide-level alignment improves accuracy relative to k-mer-based classifiers and preserves species-level resolution that is often lost in protein-based approaches. By retaining alignment coordinates and mapping reads directly to annotated genomic features, MetaTracer enables direct attribution of gene expression to specific microbial species. On simulated datasets, MetaTracer achieves high accuracy for both taxonomic and gene assignment. Applied to real dental plaque metatranscriptomic datasets, MetaTracer resolves species-specific transcriptional activity and detects reproducible differences in microbial gene expression between children with early childhood caries and healthy controls.

**Availability and implementation:** MetaTracer is implemented as a Python-based workflow wrapper (metatracer v0.1.0) that depends on the mtsv-tools core engine (v2.1.0), which is written in Rust. The required functionality is supported by the v2.1.0 release of mtsv-tools. Both packages are open source under the MIT license and are available at github.com/FofanovLab/metatracer and github.com/FofanovLab/mtsv-tools. Versioned releases are archived at Zenodo (DOI: 10.5281/zenodo.18665766 and DOI: 10.5281/zenodo.18718002). Installation is supported via Bioconda.

## Background

Metatranscriptomic sequencing is widely used to measure microbial activity by quantifying expressed genes in complex communities. Many existing analysis approaches either summarize activity without taxonomic context, profile active species using marker genes without capturing the full complement of expressed genes, or link expression at course taxonomic levels that mask species-specific differences. These limitations prevent precise attribution of functional activity to individual organisms which obscures their biological roles and limits our ability to interpret microbial community function. Overcoming these limitations to achieve species-resolved, genome-wide gene expression is a central goal of metatranscriptomic analysis.

Current metatranscriptomic workflows are fragmented, typically separating taxonomic classification from gene assignment. Taxonomic characterization is often performed using k-mer-based read classifiers (e.g. Kraken2 (Wood and Salzberg 2014; Wood, Lu and Langmead 2019)) or marker-gene profilers (e.g. MetaPhlAn (Blanco-Míguez *et al*. 2023)); however, these approaches do not retain genomic coordinates and therefore cannot directly map transcripts to genes. Functional profiling is carried out in a separate step using translated protein alignment tools (e.g., DIAMOND-based pipelines (Buchfink, Xie and Huson 2015; Buchfink, Reuter and Drost 2021) such as HUMAnN (Beghini *et al*. 2021), SAMSA (Westreich *et al*. 2016), or MEGAN (Huson *et al*. 2007)). Although protein-level searches are computationally efficient due to smaller and more tractable databases, translation collapses nucleotide-level variation, reducing species-level resolution and necessitating a separate taxonomic classification step to achieve higher resolution.

To address these limitations, we present MetaTracer, a nucleotide-level, full-alignment metatranscriptomic analysis tool that simultaneously performs high-resolution taxonomic classification and gene assignment. MetaTracer builds on our previously introduced alignment-based taxonomic classifier, MTSv (Furstenau et al. 2022), which provides accurate species-level classification with improved tolerance to sequence divergence, and fewer false positives than k-mer-based methods. To avoid the computational cost of exhaustive alignment against large reference databases, MTSv employs a taxon-aware alignment strategy that enables tens of millions of reads to be processed within a few hours. Although MTSv cannot match the speed of marker or k-mer-based tools, the key advantage is the ability to retain precise genomic alignment coordinates. This property has been leveraged in MetaTracer, enabling direct mapping of reads to annotated genomic features. Consequently, each read can be assigned simultaneously to both a species and a gene in a single pass, eliminating the need for separate taxonomic and functional pipelines.

## Methods

### Overview

MetaTracer uses a reference-based, full-alignment framework to jointly assign taxonomic and gene annotations to sequencing reads. The pipeline first constructs a metagenomic FM-index (Ferragina and Manzini 2000), previously implemented in MTSv (Furstenau *et al*. 2022), from a collection of reference genome sequences. Candidate alignment regions are identified using FM-index-based seeding and valid candidates are evaluated using full nucleotide alignment. For each accepted alignment, the corresponding taxonomic identifier, reference genome, and genomic coordinate are recorded and used to query pre-indexed gene annotation files for functional assignment.

### Reference database construction and indexing

Reference genomes are retrieved from the NCBI RefSeq database (O’Leary *et al*. 2016) using the NCBI Datasets command-line tool (v18.3.1) (O’Leary *et al*. 2024). To support accurate transcript identification, MetaTracer relies on well-annotated reference genomes from which gene coordinates can be obtained and indexed. Because alignment-based approaches are tolerant to sequence divergence, exhaustive strain-level representation is unnecessary, provided that the species pangenome is adequately represented. In this study, species were limited to a maximum of 100 genomes, resulting in 25,143 total genome accessions representing 5,940 bacterial species (see Data Availability for a complete list of assembly accessions). To improve scalability and accommodate varying computational resources, the reference database is partitioned into configurable chunks that are indexed independently. This design enables parallel index construction and alignment, allows memory requirements to be tuned by adjusting chunk size, and supports incremental updates or customization of the reference set without rebuilding the entire data structure. For this study, the reference database was divided into ten FASTA files (∼10 GB each) producing index files of approximately 35 GB each.

### Taxonomic assignment

Reads are processed independently against each index file. Each read is decomposed into fixed-length seeds, which are queried against the index to identify candidate regions meeting a user-defined seed hit threshold. Candidate regions are evaluated in descending order of seed support, and a SIMD-accelerated Smith-Waterman alignment (Zhao *et al*. 2013) is performed for each until an alignment meeting a user-defined edit distance threshold is identified. Once a passing alignment is found for a given taxon, no further alignments are evaluated for that taxon, reducing redundant computation when multiple reference genomes are present. Results from all indices are merged into a single report containing taxonomic assignments, genome identifiers, alignment offsets, and edit distances. Full nucleotide-level alignment enables many reads to be uniquely assigned; for ambiguous reads, taxa with higher edit distances or low sample-level abundance can be filtered, and any unresolved reads are reported as multi-taxon assignments.

### Gene assignment

Once taxonomic assignments are determined, alignment coordinates for the selected taxon are used to query pre-indexed GFF annotation files and identify overlapping coding regions. Reads overlapping annotated genes are assigned to the corresponding gene identifiers along with associated metadata. The corresponding protein sequences are retrieved from the reference protein FASTA file, enabling downstream homology-based functional annotation using external tools like eggNOG (Huerta-Cepas et al. 2019; Cantalapiedra et al. 2021). Read-level assignments can then be aggregated to generate count matrices that are compatible with external differential expression and pathway-level analysis tools.

### Read simulation

Simulated metatranscriptomic datasets were generated to assess taxonomic and gene-level assignment accuracy. Community composition and relative abundances were modeled after the CAMI2 human oral microbiome benchmark datasets (Fritz *et al*. 2019; Meyer *et al*. 2022). For each taxon, CDSs from a single representative genome were used, and mutations were introduced using the Mason simulator v2.0.12 (Holtgrewe 2010). For each of the 10 samples, 10 million 150bp single-end Illumina reads were simulated. Quality control was performed using fastp (v0.23.4) (Chen 2023). Simulated reads, scripts, and benchmarking data have been deposited in a public repository (see Data Availability)

### Comparison of taxonomic assignment performance with Kraken2

To benchmark taxonomic assignment performance, MetaTracer was compared against Kraken2 v2.1.3 (Wood and Salzberg 2014; Wood, Lu and Langmead 2019), a widely used nucleotide-based taxonomic classifier. For direct comparison, analyses were based on read-level classification rather than the abundance estimates produced by Kraken2 or Bracken (Lu *et al*. 2017). The Kraken2 database was constructed using the same set of reference genomes used for MetaTracer, and default parameters were used for classification.

### Analysis of real metatranscriptomic data from pediatric dental plaque samples

MetaTracer was evaluated using real metatranscriptomic datasets generated from dental plaque samples collected from children with (n=15) and without (n=19) early childhood caries (ECC). Samples from ECC children were collected from caries lesions, while samples from caries-free children were obtained from healthy tooth surfaces. Total RNA was extracted from plaque material, subjected to rRNA depletion using QIAGEN FastSelect rRNA 5S/16S/23S kits (Qiagen, Hilden, Germany), and sequenced using Illumina 2×150bp paired-end RNA sequencing. A complete description of the methods have been reported previously (Hsu *et al*. 2024) and the sequencing reads have been deposited in a public repository (see Data Availability).

### Differential expression and pathway enrichment analysis

From the dental plaque metatranscriptomic dataset, a taxon-resolved gene count matrix was constructed by aggregating read counts at the gene level within each taxon. To reduce noise from low-abundance features, we filtered those that did not have 5 or more reads in 3 or more samples per group (caries-free vs. ECC). Differential expression (DE) analysis between ECC and caries-free children was performed using DESeq2 v1.50.2 (Love, Huber and Anders 2014). Features were considered differentially abundant if they met a Benjamini-Hochberg adjusted p-value (FDR) < 0.05 and an absolute log_2_ fold change ≥ 1.

## Results

### Taxonomic assignment accuracy

Across 10 simulated datasets, MetaTracer assigned an average of 99.99% of reads, 99.86% (IQR: 99.86– 99.88) of which were assigned to the true taxon. Among correctly assigned reads, 90.5% (IQR: 89.28–92.50) were unambiguously assigned to a single species. For comparison, Kraken2 assigned a similar proportion of reads (99.99%) with 99.5% (IQR: 99.52–99.61) assigned correctly, and 88.0% (IQR: 86.87–90.06) assigned at the species level. Consistent with our previous publication (Furstenau *et al*. 2022), Kraken2 showed reduced performance for reads that diverged from reference sequences and introduced substantially more false-positive species-level assignments, with an average of 446 unexpected species per simulation compared to 41 for MetaTracer, and approximately six-fold more reads assigned to these spurious taxa.

### Gene assignment accuracy

An average of 95.98% (IQR: 95.68–96.34) of taxonomically assigned reads overlapped annotated coding regions and could be linked to gene features. Because gene identifiers can vary across genomes, eggNOG v2.1.12 (Huerta-Cepas *et al*. 2019; Cantalapiedra *et al*. 2021) orthologous groups (OG) assignments were used to standardize gene categories when assessing accuracy. On average, 97.86% (IQR: 97.38–98.09) of reads were assigned to the same OG as the sequence they were simulated from, and of those 88.98% (IQR: 87.78–91.17) were previously assigned at the species level. In comparison, the eggNOG output indicated that over 99% of amino acid sequences in the sample could only be resolved to the genus level or higher, demonstrating that MetaTracer’s nucleotide-level alignment strategy provides substantially greater species-level resolution relative to protein-based homology approaches.

### Performance with early childhood caries data

Species-resolved gene expression counts were generated using MetaTracer and used in differential expression analysis between children with ECC and caries-free controls. A total of 9,445 OGs were significantly differentially expressed (DE) (adjusted p<0.05, |log_2_FC|>=1). The most transcriptionally active organisms included well established contributors to ECC, and the upregulated functional categories were consistent with known cariogenic activities (Loesche 1986a, 1986b; Takahashi and Nyvad 2011; Edlund *et al*. 2015), including carbohydrate metabolism, biofilm formation, acid production, and stress tolerance (Figure 1A). Figure 1B illustrates the importance of species-level resolution for interpreting transcriptional dynamics in complex microbial communities. When transcripts were aggregated at the genus level—approximating the resolution typically achieved by protein-sequence-based approaches—most differential expression signals were lost or reduced, as reads from closely related species were averaged within a single taxonomic unit. In communities where multiple related species co-occur, this aggregation obscures opposing or heterogeneous transcriptional responses, masking biologically meaningful patterns. By providing accurate species-level gene assignments, MetaTracer preserves these signals and allows distinct functional activities to be disentangled within complex microbial communities.

**Figure 1.**
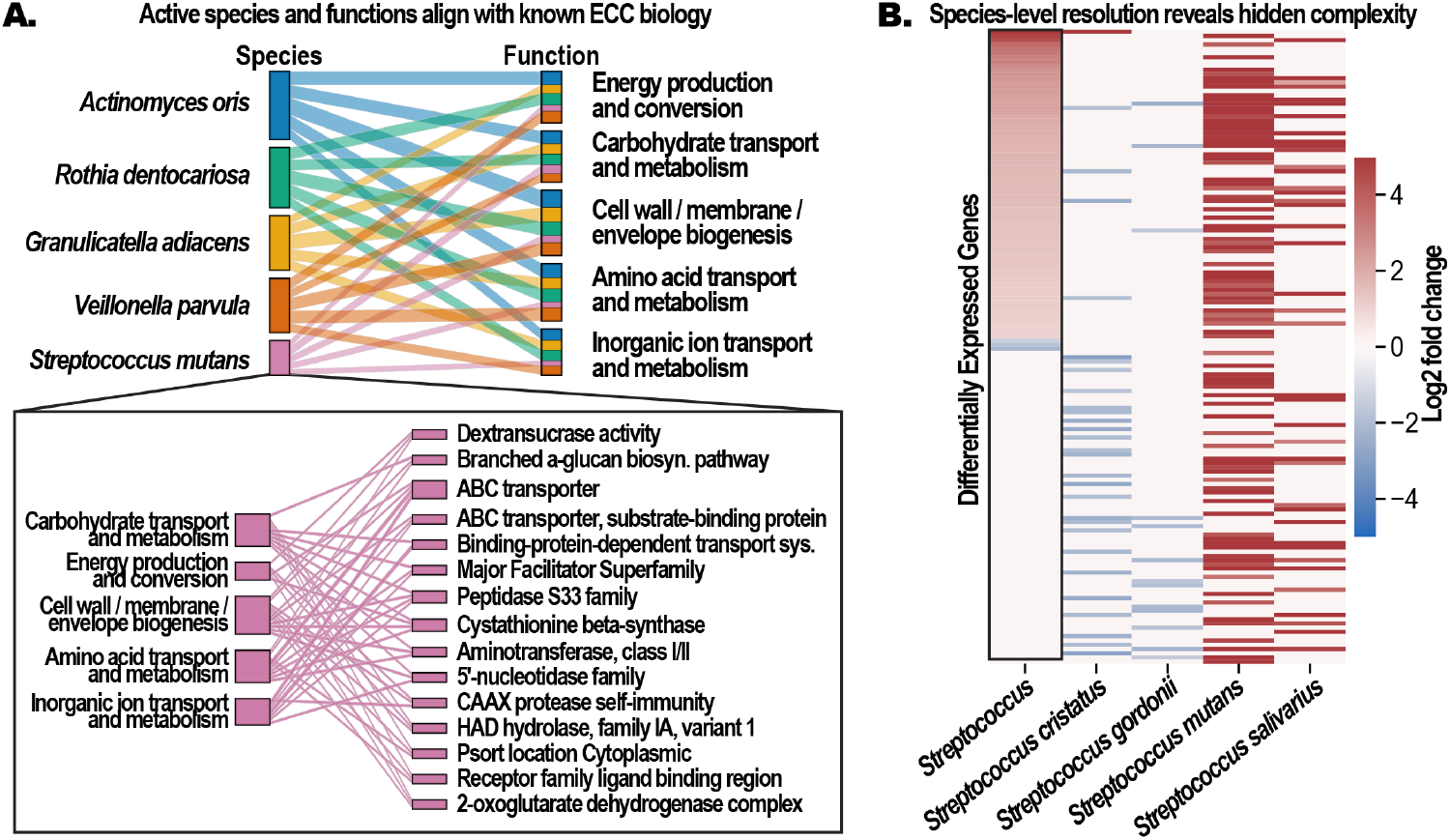
Species-resolved metatranscriptomic profiling recapitulates known caries-associated organisms and functions while revealing functional heterogeneity obscured by genus-level aggregation. A) Sankey diagram summarizing the five most transcriptionally active bacterial species identified by MetaTracer in ECC samples and the functional categories associated with their significantly upregulated genes. B) Heatmap comparing genus- and species-level DE within *Streptococcus*. The leftmost column shows log2 fold changes after genus-level aggregation, while the remaining columns show species-resolved patterns for four species. Species-level patterns reveal heterogeneous activity among co-occurring species that are attenuated or missing from the aggregated result.

## Conclusions

MetaTracer is a metatranscriptomic analysis tool designed for complex microbial communities that simultaneously performs taxonomic classification and gene assignment within a single analytic pass. Across simulated and real datasets, the method achieved high taxonomic and gene-assignment accuracy, reduced false-positive classifications relative to k-mer-based classifiers, and maintained species-level functional resolution that is often lost with protein-based workflows. Applied to real oral microbiome samples, MetaTracer recapitulated established cariogenic taxa and pathways and revealed species-specific differential expression patterns that were obscured when transcripts were aggregated at higher taxonomic levels. These findings demonstrate that retaining species-level gene assignments improves the interpretation of microbial activity and enables functional differences among closely related organisms to be distinguished, providing clearer insight into community organization, interactions, and ecological niche structure.

## Data Availability

Oral microbiome sequencing reads have been deposited in the NCBI Sequence Read Archive under BioProject accession PRJNA1024640. Simulated reads, data, and analysis scripts are available at Figshare DOI:10.6084/m9.figshare.31245190.

## Acknowledgements

Computational resources and support were provided by the Northern Arizona University Advanced Research Computing (ARC) group and the Monsoon High Performance Computing cluster.

## References

Beghini F, McIver LJ, Blanco-Míguez A et al. Integrating taxonomic, functional, and strain-level profiling of diverse microbial communities with bioBakery 3. Turnbaugh P, Franco E, Brown CT (eds.). eLife 2021;10:e65088.

Blanco-Míguez A, Beghini F, Cumbo F et al. Extending and improving metagenomic taxonomic profiling with uncharacterized species using MetaPhlAn 4. Nat Biotechnol 2023;41:1633–44.

Buchfink B, Reuter K, Drost H-G. Sensitive protein alignments at tree-of-life scale using DIAMOND. Nat Methods 2021;18:366–8.

Buchfink B, Xie C, Huson DH. Fast and sensitive protein alignment using DIAMOND. Nat Methods 2015;12:59–60.

Cantalapiedra CP, Hernández-Plaza A, Letunic I et al. eggNOG-mapper v2: Functional Annotation, Orthology Assignments, and Domain Prediction at the Metagenomic Scale. Mol Biol Evol 2021;38:5825–9.

Chen S. Ultrafast one-pass FASTQ data preprocessing, quality control, and deduplication using fastp. Imeta 2023;2:e107.

Edlund A, Yang Y, Yooseph S et al. Meta-omics uncover temporal regulation of pathways across oral microbiome genera during in vitro sugar metabolism. ISME J 2015;9:2605–19.

Ferragina P, Manzini G. Opportunistic data structures with applications. 41st Annu Symp Found Comput Sci Proc 2000:390–8.

Fritz A, Lesker T, Bremges A et al. CAMI 2 Multisample Benchmark Dataset of the Human Microbiome Project. 2019, DOI: 10.4126/FRL01-006425518.

Furstenau TN, Schneider T, Shaffer I et al. MTSv: rapid alignment-based taxonomic classification and high-confidence metagenomic analysis. PeerJ 2022;10:e14292.

Holtgrewe M. Mason: A Read Simulator for Second Generation Sequencing Data. Tech Rep FU Berl 2010.

Hsu K-LC, Shaffer IN, Furstenau TN et al. Ethnicity-specific microbiome in early childhood caries: from the functional perspectives of oral biofilm. Res Sq 2024, DOI: 10.21203/rs.3.rs-5153245/v1.

Huerta-Cepas J, Szklarczyk D, Heller D et al. eggNOG 5.0: a hierarchical, functionally and phylogenetically annotated orthology resource based on 5090 organisms and 2502 viruses. Nucleic Acids Res 2019;47:D309–14.

Huson DH, Auch AF, Qi J et al. MEGAN analysis of metagenomic data. Genome Res 2007;17:377–86.

Loesche WJ. Role of Streptococcus mutans in human dental decay. Microbiol Rev 1986a;50:353–80.

Loesche WJ. The identification of bacteria associated with periodontal disease and dental caries by enzymatic methods. Oral Microbiol Immunol 1986b;1:65–72.

Love MI, Huber W, Anders S. Moderated estimation of fold change and dispersion for RNA-seq data with DESeq2. Genome Biol 2014;15:550.

Lu J, Breitwieser FP, Thielen P et al. Bracken: estimating species abundance in metagenomics data. PeerJ Comput Sci 2017;3:e104.

Meyer F, Fritz A, Deng Z-L et al. Critical Assessment of Metagenome Interpretation: the second round of challenges. Nat Methods 2022;19:429–40.

O’Leary NA, Cox E, Holmes JB et al. Exploring and retrieving sequence and metadata for species across the tree of life with NCBI Datasets. Sci Data 2024;11:732.

O’Leary NA, Wright MW, Brister JR et al. Reference sequence (RefSeq) database at NCBI: Current status, taxonomic expansion, and functional annotation. Nucleic Acids Res 2016;44:D733–45.

Takahashi N, Nyvad B. The role of bacteria in the caries process: ecological perspectives. J Dent Res 2011;90:294–303.

Westreich ST, Korf I, Mills DA et al. SAMSA: a comprehensive metatranscriptome analysis pipeline. BMC Bioinformatics 2016;17:399.

Wood DE, Lu J, Langmead B. Improved metagenomic analysis with Kraken 2. Genome Biol 2019;20:257.

Wood DE, Salzberg SL. Kraken: ultrafast metagenomic sequence classification using exact alignments. Genome Biol 2014;15:R46.

Zhao M, Lee WP, Garrison EP et al. SSW library: an SIMD Smith-Waterman C/C++ library for use in genomic applications. PLoS One 2013;8:e82138.

